# The origin and maintenance of microbial symbionts in *Drosophila* larvae

**DOI:** 10.1101/2020.09.12.294868

**Authors:** Robin Guilhot, Auxane Lagmairi, Laure Olazcuaga, Anne Xuéreb, Simon Fellous

## Abstract

Little is known on the origin and maintenance of symbionts associated with *Drosophila* larvae *in natura*, which restricts the understanding of *Drosophila*-extracellular microorganism symbiosis in the light of evolution. Here, we studied the origin and maintenance of symbionts of *Drosophila* larvae under ecologically realistic conditions, to our knowledge for the first time, using yeast and bacterial isolates and two *Drosophila* species: the model organism *D. melanogaster* and the invasive pest *D. suzukii*. We discovered that *Drosophila* females and males both transmit yeast and bacteria symbionts to larvae. In addition, several symbiotic yeasts initially associated with larvae were conserved throughout host life cycle and transmitted to offspring. Our results suggest that stable associations of *Drosophila* flies with bacteria and yeasts may exist *in natura* and constitute a step forward in the understanding of wild *Drosophila*-microorganism symbioses.

## Context

The origin of microbial symbionts of eukaryotes influences the evolution of symbiosis. Microbial symbionts can be acquired from parents (i.e. vertically transmitted symbionts) (Funkhouser & Bordenstein 2013), from unrelated hosts (i.e. horizontally transmitted symbionts) (Gonella *et al*. 2012), a mix of both (i.e. mixed-mode transmitted symbionts) (Ebert 2013; Quigley *et al*. 2018) or from the environment (Kikuchi *et al*. 2007). Theory predicts that symbionts that persist between host life stages and host generations are more likely to initiate stable mutualistic relationships compared to symbionts acquired from the host environment (Antonovics *et al*. 2017; Bright & Bulgheresi 2010; Fisher *et al*. 2017; Gerardo & Hurst 2017; Lipsitch *et al*. 1996; Sachs *et al*. 2004; Shapiro & Turner 2014). Understanding the evolution of host-microbe symbiosis is therefore only possible when means of host-microbe association are properly documented.

In *Drosophila* flies, numerous studies conducted under laboratory conditions investigated the origin of extracellular microbial symbionts associated with larvae and the persistence of larval symbionts throughout host life cycle (Bakula 1969; Becher *et al*. 2012; Pais *et al*. 2018; Téfit *et al*. 2018). However, little is known on the origin and maintenance of symbionts associated with *Drosophila* larvae *in natura*, which restricts the understanding of *Drosophila*-extracellular microorganism symbiosis in the light of evolution. We explored these phenomena under ecologically realistic conditions, to our knowledge for the first time, using - mainly wild - yeast and bacterial isolates and two *Drosophila* species of major interest: the model organism *D. melanogaster* and the invasive pest *D. suzukii*.

## Results and Discussion

### *Drosophila* females transmit extracellular symbionts to their offspring

Previous reports showed *D. melanogaster* maternal transmission of yeasts and bacteria in laboratory conditions (Bakula 1969; Becher *et al*. 2012; Rohlfs and Hoffmeister 2005; Spencer *et al*. 1992; Téfit *et al*. 2018). We hypothesized that *Drosophila* mothers may transmit their microbial symbionts to larvae in a context where other microorganisms are present on the oviposition substrate. We also predicted that *D. suzukii* maternal transmission may be more frequent that of *D. melanogaster* because *D. suzukii* females typically lay their eggs on unwounded, ripening fruits poorly colonized by microorganisms (Lewis & Hamby 2019). *D. suzukii* eggs are inserted in fruit flesh thanks to females’ serrated ovipositors. As a result, the newly emerged larvae may primarily recruit microbial symbionts deposited by the mother. By contrast, *D. melanogaster* females lay their eggs on fruit wounds and rotten fruits already colonized by a variety of microorganisms (data not shown, will be available in the next version of this work). To test these predictions, we used mature females of one *D. suzukii* population and one *D. melanogaster* population and six microbial symbionts (see Materials and Methods for details on their choice). The same microbial strains were used for all the experiments presented in our study. Briefly, individual mated female associated with an artificial microbial community composed of one bacterium and one yeast strain were offered to oviposit on a blueberry which surface had been inoculated with a different microbial community (i.e. another bacterium and another yeast) (Figure 1A). For *D. melanogaster* assays the berry was slightly wounded while kept unwounded for *D. suzukii* assays. Five days after fruit exposure, numerous berries contained larvae associated with female microbial symbionts, fruit-surface microorganism or both (Figure 1B).

**Figure 1.**
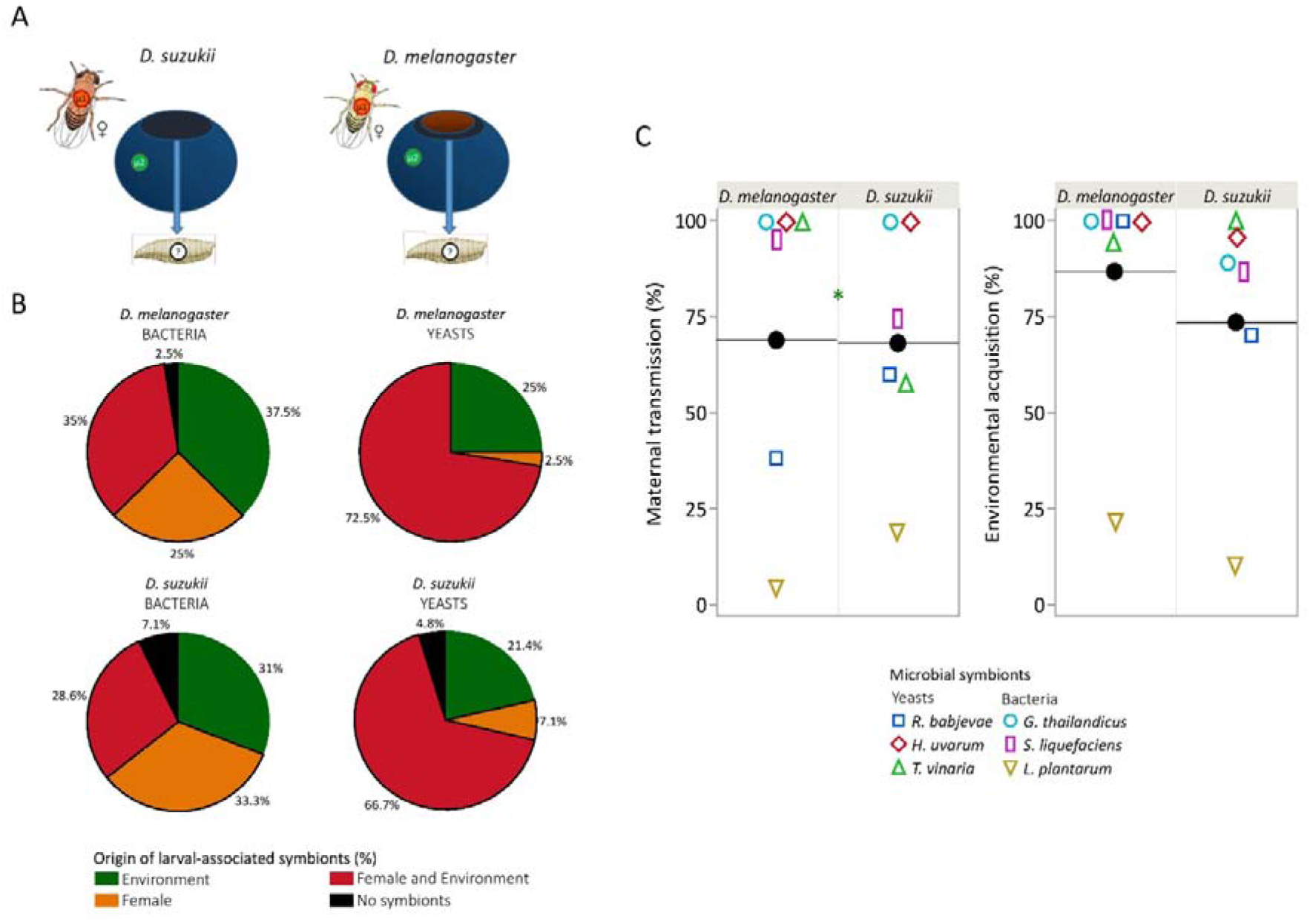
*Drosophila* larvae associate with maternal symbionts and environmental symbionts. (A) Experimental design. Three different microbial communities (pi,) composed of a yeast and a bacterium species were permuted between flies and fruits. n = 40 *D. melanogaster* experimental units; n = 42 *D. suzukii* experimental units. (B) *Drosophila* larvae frequently harbored maternal symbionts and those already present on fruit skin. (C) Maternal transmission and environmental acquisition rates (% of larvae pools). The black dot symbolizes the general mean (i.e. independently of the microbial symbiont) and the open symbols the proportion for each of the six microorganisms tested.

Contrary to our expectations, maternal transmission was no greater in *D. suzukii* than in *D. melanogaster*. However, symbiont identity affected both maternal transmission and environmental acquisition (Table S1). One yeast strain, *Trigonopsis vinaria* isolated from *D. suzukii* ovaries was transmitted significantly more from *D. melanogaster* than from *D. suzukii* females (χ^2^ = 5.25, df = 1, p = 0.0220) (Figure 1C, Table S1). Symbiont transmission differed whether they were in females or on fruit, which suggests acquisition of maternal symbionts by offspring is controlled by interactions between females and symbionts rather than symbiont’s sheer properties. Our work indicates *D. suzukii* and *D. melanogaster* maternal transmission of extracellular symbionts may be frequent in field conditions.

### Male transmission of microbial symbionts

How microorganisms reach fruit skin, where they are recruited by *Drosophila* larvae, is unclear. Insects, such as wasps, participate to baker’s yeast spread at the landscape level (Stefanini *et al*. 2012). Field observations showed *Drosophila* males often sit on fruit, a behavior that we also witnessed in lab macrocosms (SM2). We hence hypothesized *Drosophila* males may deposit their symbionts on fruit surface and therefore contribute to the larval microbiota. *D. melanogaster* males can be territorial, can form leks and defend oviposition sites (Drapeau *et al*. 2011; Hoffmann and Cacoyianni 1990). Because *D. melanogaster* males are present on fruit wounds (i.e. oviposition sites), where microorganisms could grow better than on fruit skin, we predicted greater male transmission from *D. melanogaster* than from *D. suzukii*. In a new experiment we tested whether *Drosophila* males actually transmitted their microbial symbionts to offspring of conspecific females (Figure 2A). Individual males were given single blueberries for 24 h until single females were added for another 24 h. As before, males, females and fruits were all associated with different microbial communities.

**Figure 2.**
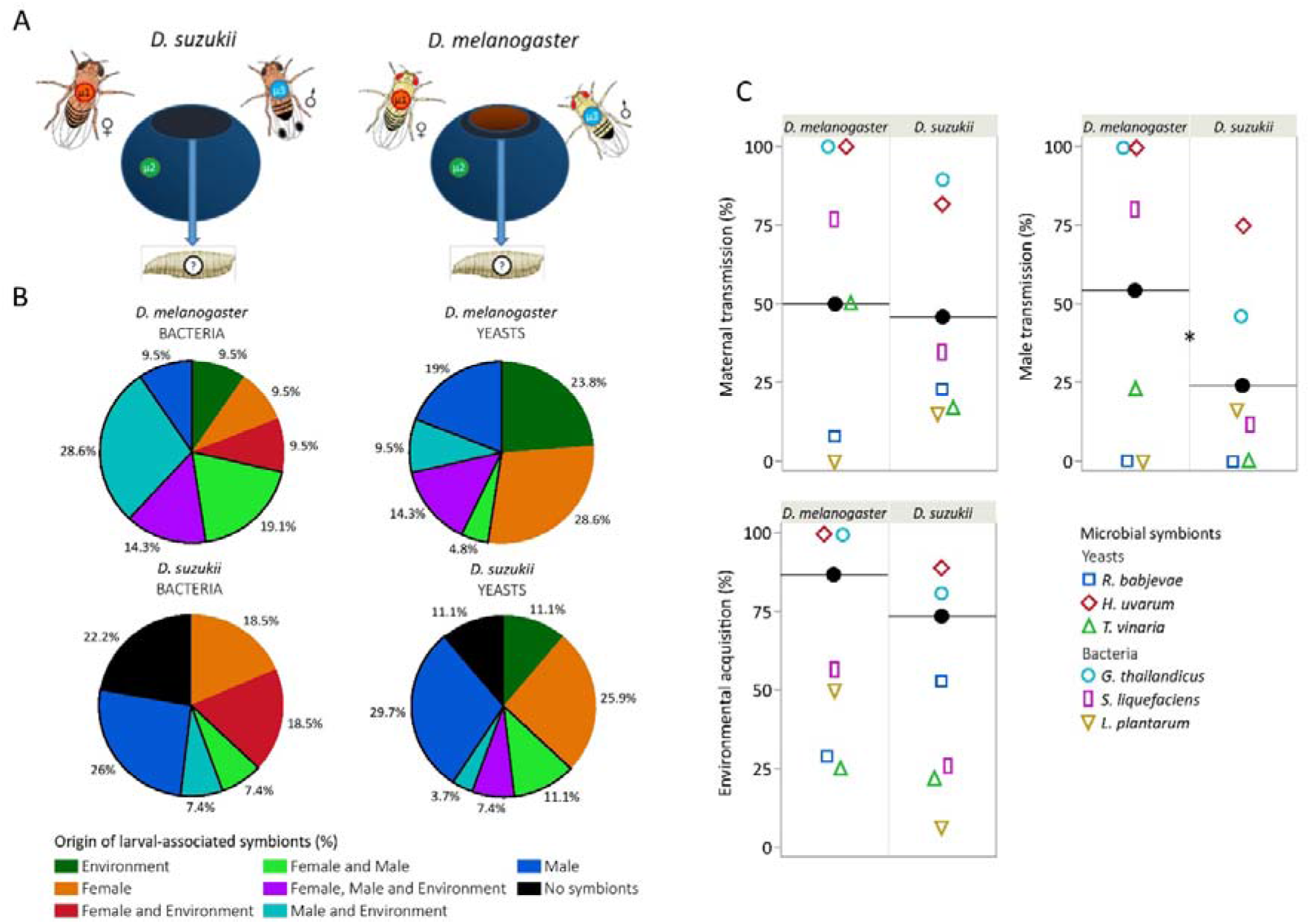
*Drosophila* larvae associate with male symbionts, maternal symbionts and environmental symbionts. (A) Experimental design. Three different microbial communities (μ_i_) composed of a yeast and a bacterium species were permuted between flies and fruit, n = 21 *D. melanogaster* experimental units; n = 27 *D. suzukii* experimental units. (B) *Drosophila* larvae frequently harbored symbionts of both male and female as well as those already present on fruit skin. (C) Proportion of cases where larvae contained male, female and fruit symbionts (% of larval pools). The black dot symbolizes the general mean (i.e. independently of the microbial symbiont) and the open symbols the proportion for each of the 6 microorganisms tested.

Male transmission to larvae was pervasive and twice more frequent for *D. melanogaster* (c. 50% of fruits) than *D. suzukii* (c. 25%) (Figures 2B and 2C). The transmission by males of the microorganisms widely depended on strain identity. For example, the yeast *H. uvarum* was always transmitted by *D. melanogaster* males while the yeast *R. babjevae* was never found in larvae. Female transmission was slightly lower than that of in the first experiment, with different behaviors of the microbial strains (Figure 2C, Table S3). The transmission potential of the symbiont strains appeared different in males and females and among experiments in females suggesting this aspect of strain biology is very context-sensitive.

How did males transmit their symbionts? We recorded the time they spent on oviposition areas but this variable did not correlate significantly with the transmission of their symbionts to larvae (Table S3). Male transmission is therefore not determined by the amount of microbial cells they shed on oviposition sites. Similarly, we recorded whether males and females mated during the experiment. These events were rare (n = 7/21 observations for *D. melanogaster* and n = 2/27 for *D. suzukii*) and did not influence significantly male transmission (Table S3). This suggests that male transmission of symbionts to larvae did not depend from male presence on oviposition sites and did not clearly involve sexual transmission to females (Miest and Bloch-Qazi 2008; Rohlfs and Hoffmeister 2005; Starmer, Peris and Fontdevila 1988). Independent of the mechanisms, the transmission of symbionts by *Drosophila* males may have consequences for the evolution of symbionts effects on males. Microorganisms may change male characters so as to favor their transmission. Because symbiont transmission is not contingent upon male reproduction (i.e. no male vertical transmission), selection may not select against symbiont costs to male fitness (Sachs 2004; Ebert 2013). However, the largest *D. melanogaster* males would be most likely to successfully defend oviposition sites (Hoffmann 1987). Therefore, symbionts of male larvae would be selected for beneficial effects on their development, assuming that extracellular symbionts of larvae remain associated with their hosts after metamorphosis and until they reproduce.

### Larval yeast symbionts maintain through the entire life cycle and transmit to the progeny

Do extracellular symbionts of *Drosophila* larvae maintain until adult life? Several studies have shown symbionts of larvae can be found in adults (i.e. transstadial transmission, maintenance through metamorphosis) (Bakula 1969; Duneau and Lazzaro 2018; Ridley *et al*. 2012; Starmer, Peris and Fontdevila 1988). However, in most experiments larvae and adults shared the same containers hence permitting indirect, environmental transmission (but see Bakula 1969). In the field, *Drosophila* last-instar larvae mainly pupate outside the larval environment, usually in soil (Reaume & Sokolowski 2006; Woltz & Lee 2017). This behavior was mimicked in a new experiment where *Drosophila* larvae were associated with one of three yeast strains, newly formed pupae isolated in independent containers and adult microbial content assayed shortly after emergence (Figure 3A).

**Figure 3.**
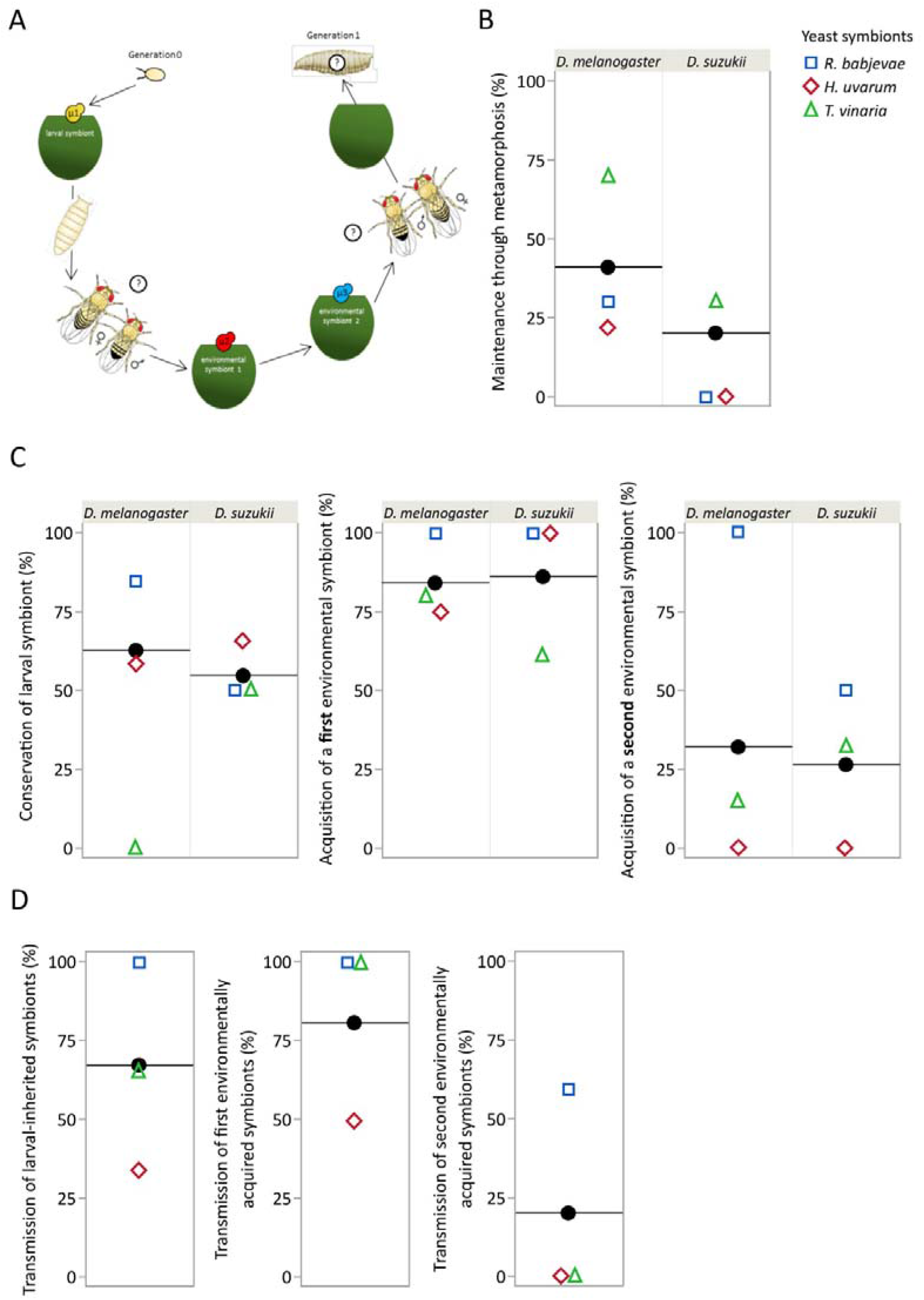
Larval yeast symbionts maintain through *Drosophila* stages and generations. (A) Experimental design. μ means microbial community. (B) Maintenance of symbionts through metamorphosis. (C) Maintenance of larval symbionts and acquisition of environmental symbionts in adults. (D) Transmission of the different adult symbionts to a new fly generation. The black dot symbolizes the general mean (i.e. independently of the microbial symbiont).

Almost a quarter and a half of young *D. suzukii and D. melanogaster* adults tested positive for larval symbionts (cell numbers where however usually low). *Trigonopsis vinaria* yeast, isolated from *D. suzukii* ovaries, best maintained through host metamorphosis while our strain of *Hanseniaspora uvarum*, a species frequently found in wild Drosophilids, exhibited poor transstadial transmission (χ^2^ = 0.0188, df = 2, p = 0.0188) (Figure 3B). Fly species and adult sex had marginally non-significant influences on yeast transstadial maintenance (for both, χ^2^ = 3.70, df = 1, p = 0.0544) (Figure 3B). Overall, yeast symbionts of larvae maintained until adult emergence, but what would become of them remained to be determined.

Numerous laboratory experiments indicate adult symbiotic community mirrors that of their surrounding environment due to the constant replacement of gut microbial communities during feeding (Blum *et al*. 2013; Ma & Leulier 2018). However, other work shows adult association with some nutritional symbionts, in particular those recently isolated from the field, may be stable (Pais *et al*. 2018; Obadia *et al*. 2017). We continued the previous experiment in order to determine whether yeast present in young adults, and acquired at the larval stage, maintained through life and until the next generation (Figure 3A). Freshly emerged adults associated with one of three yeast strains were maintained for five days with a halved grape berry inoculated with a second yeast strain (i.e. first environmental symbiont in Figure 3A) and another two days with berries inoculated with a third strain (i.e. third environmental symbiont). These adults were then offered a surface-sterilized berry to oviposit. The microbial content of adults at the time of oviposition and that of F1 larvae was assayed.

Unexpectedly, yeast symbionts of larval origin largely maintained despite a one-week exposure to two successive sources of environmental yeasts (Figure 3C, Table S4). The second environmental yeast was less frequent in adults compared to the first (Figure 3C). Even if the time of adult exposure to symbionts may affects host acquisition (as in Obadia *et al*. 2017), it should be mentioned we did not monitor microbial development in the grape berries the flies were exposed to for five and two days, respectively. It is therefore possible flies inoculated them with the strains they harbored, hence favoring the multiplication of larval and first environmental symbionts in our microcosms and their subsequent re-inoculation to adults. Incidentally, it could explain the greater prevalence of larval yeast in old adults than in young adults (compare Figures 3B and 3C). Nonetheless, the majority of *D. melanogaster* larvae of the following generation bore symbionts their parents were first exposed to at the larval stage (proportion of larvae with the larval symbionts of their parents: 0.69 (95% CI [0.44, 0.86])) (Figure 3D). This experiment shows that, in field-realistic conditions, symbiotic yeasts associated with *D. melanogaster* and *D. suzukii* larvae are conserved in young adults despite metamorphosis and illustrate symbiont persistence throughout host life cycle until they are transmitted to a new generation.

## Conclusion

We discovered that *Drosophila* females and males both transmit their extracellular symbionts to larvae. Several symbiotic yeasts initially associated with larvae were conserved throughout host life cycle and transmitted to offspring. Our results, mainly obtained with microorganisms freshly isolated in the wild, suggest that stable associations of *Drosophila* flies with bacteria and yeasts may exist *in natura*. As our results were obtained under ecologically realistic conditions, they may therefore constitute a tangible step forward in the understanding of wild *Drosophila* - microorganism symbioses.

A major issue in the recent *Drosophila* literature is to determine how exactly microbial symbionts maintain association with the host. Most studies conclude *Drosophila* microbial symbionts do not maintain durably in the host. Symbionts would be continuously inoculated by the host to the substrate where they multiply, reacquired from the environment via a ‘farming’ mechanism but rarely conserved in absence of intake during feeding. However, this phenomenon has been described using laboratory strains of *Drosophila* and symbionts under typical laboratory conditions (Blum *et al*. 2013; Storelli *et al*. 2018). Along these lines, several studies show extracellular symbionts found in arthropods reflect the microbial communities they encounter in their diet (Kennedy *et al*. 2020; Moran *et al*. 2019). By contrast, evidence of the existence of resident extracellular symbionts of *Drosophila* accumulates. In *D. melanogaster* adults, two recent independent studies show that wild isolates of the bacteria *Lactobacillus plantarum* and *Acetobacter thailandicus* may durably colonize the first gut region of the host (crop, crop duct and proventriculus) independently of the ingestion of other symbionts under laboratory conditions (Pais *et al*. 2018; Obadia *et al*. 2017). In the wild, such resident symbionts may durably persist in host individuals and populations. Our study was not designed to investigate how and why extracellular *Drosophila* symbionts persist in or get lost by adult hosts. However, we found symbiont maintain throughout metamorphosis, a phenomenon that was poorly studied with wild strains in fruit (Bakula 1969; Ridley *et al*. 2012; Starmer *et al*. 1988). Differences among yeasts strains in terms of maintenance and transmission may relate to where they locate in the host and therefore how we sampled them. Indeed, the yeast *Trigonopsis vinaria* we isolated from *Drosophila* ovaries best maintained throughout metamorphosis (Figure 4B). However, *Hanseniaspora uvarum*, a species frequent on the surface of fruit (Morais *et al*. 1995), that strongly attracts Drosophila adults and is often found associated with them, always transmitted well from adults to larvae (Figures 2 and 3). If it is not possible to generalize with a handful of microbial strains, the data suggests wild symbionts vary in their strategies of host-mediated dispersal (Jacob *et al*. 2019). Most yeast species rely on insect vectors for dispersal (Kurtzman *et al*. 2011), some may be better at attracting adults, others at transmitting among life stages or to offspring. Recent literature debates whether yeast coevolve with flies on the basis that the volatiles they produce have other functions than just to attract flies (Günther *et al*. 2019; Koerte *et al*. 2020). The contingency of each other’s fitness due to yeast maintenance during metamorphosis and transmission from adults to larvae constitutes another coevolutionary paradigm. One where symbiotic associations are not solely driven by partner choice but also by co-transmission (Sachs *et al*. 2004).When host and symbiont fitnesses correlate positively selection favors mutualistic interactions (Ebert 2013; Lipsitch *et al*. 1996; Sachs *et al*. 2004). In *Drosophila*, benevolent effects of extracellular symbionts may amount to better provisioning of nutriments (Ankrah & Douglas 2018) or host protection against pathogens (Johnston & Rolff 2015). Future research will tell whether yeast - and symbiotic bacteria - harbor adaptations favoring long-term associations with hosts and maximize their own fitness by mutualistic influence on their host.

**Figure 4.**
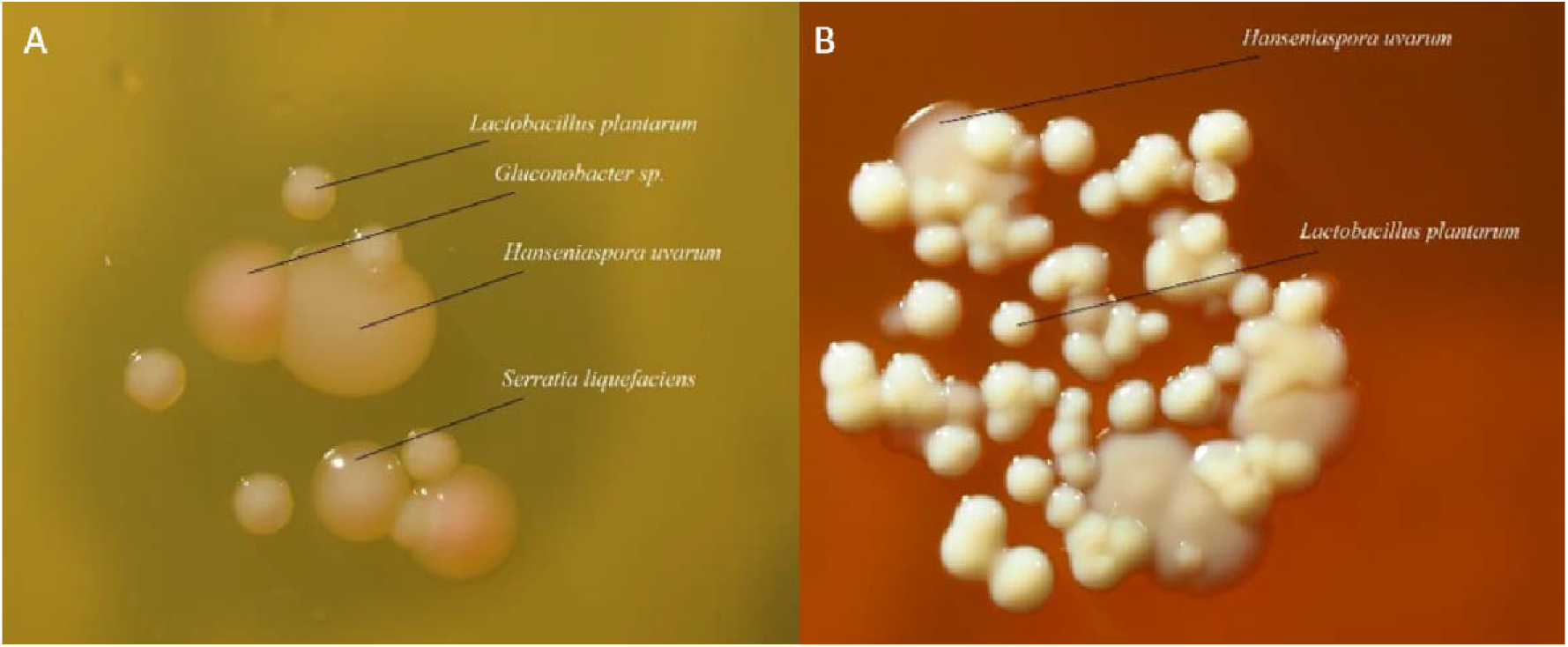
Colony morphology as a tool for discriminating mixed microbial isolates. (A) MRS agar plate (incubated at 30°C) allowed to distinguish colonies of four microbial isolates. (B) Mannitol agar plate (incubated at 24°C) allowed to distinguish colonies of two microbial isolates.

Symbiont persistence has broad consequences for the eco-evolutionary dynamics of host and symbionts in heterogeneous environments. The maintenance of symbionts over days or generations enables their participation to host adaptation to local conditions. In return benevolent symbionts may benefit improved dispersal to new resource patches. For that matter, orchards, shrubs and cities where *Drosophila* and their symbionts may be encountered resembles the very definition of meta-populations: fruits are ephemeral patches of finite resources from which it is necessary to disperse to survive in the long run. Incidentally, understanding how hosts acquire and transmit non-obligatory symbionts, such as the bacteria and yeast we studied here, helps with a major challenge for the years to come. The ecological and evolutionary dynamics of most microorganisms in space and time remains obscure, in particular in structured, complex environments (Dudaniec & Tesson 2016). Empirical study of opportunistic symbionts in natural conditions or with field-realistic microcosms will shed light on some of this mystery.

## Materials and Methods

### *Drosophila* stocks and microbial symbionts

We used two populations of *Drosophila melanogaster* (Population A, founded from OregonR individuals furnished by colleagues, and Population B, founded from wild individuals collected in the late 2017 around Montpellier, Southern France) and two populations of *D. suzukii* (Population A, founded from wild individuals collected in the early 2018 around Avignon, Southern France, and Population B, founded from wild individuals collected in 2013 in Gaujac, Southern France).

We used six microbial symbionts in this study. Five were isolated from wild flies and fruits in the late 2017. The yeasts *Hanseniaspora uvarum* MN684824, *Trigonopsis vinaria* MN684816 and *Rhodotorula babjevae* MN684819 were isolated from female *D. melanogaster* feces, *D. suzukii* ovaries and infested organic grape berries, respectively. The bacteria *Serratia liquefaciens* and *Gluconobacter thailandicus* were respectively isolated from *D. suzukii* ovaries and organic grape berries. The sixth microorganism was a laboratory isolate of the bacterium *Lactobacillus plantarum* which is widely used in bacteria – *Drosophila* studies (Ryu *et al*. 2008). Colonies of these six isolates were distinguished according to their morphology (e.g. Figure 4). More details about these isolates (their choice, their properties, the method to distinguish them) will be given in the next version of this work.

### Origin of larval microbiota

The experiments were conducted on sterile vials with conventional blueberries and gnotobiotic *Drosophila* adults, i.e. associated with particular microbial symbionts.

Gnotobiotic adults were created by inoculating axenic larvae or adults (i.e. free of extracellular symbionts here) with overnight grown microbial symbionts (MRS 30°C for *L. plantarum*, Mannitol 24°C for other bacteria, YPD 24°C for yeasts). The axenic colonies of *Drosophila* were founded with axenic eggs obtained from conventionally reared populations using a method slightly adapted from Koyle and colleagues (2016). Briefly, this method consists of removing the chorion, the outer envelope of the egg that contains extracellular microbial symbionts. The axenic colonies were maintained on sterile banana medium (water, banana, sugar, dead yeast and agar).

Blueberries were always disposed with peduncle insertion upwards. This particular zone of the berry was identified as a preferential oviposition site for *D. suzukii* females (Figure 5). To allow oviposition of *D. melanogaster* females, this zone was finely wounded using a pipette tip. All blueberries used in the experiments were surface-sterilized following the protocol of Behar and colleagues (2008). For the two main experiments of this section, surface-sterilized blueberries were artificially associated with microbial symbionts. To this aim, berries were immersed in microbial suspensions (overnight microbial culture diluted in PBS (Phosphate Buffered Saline)) and dried 18 h after a 2 min 30 vortexing.

**Figure 5.**
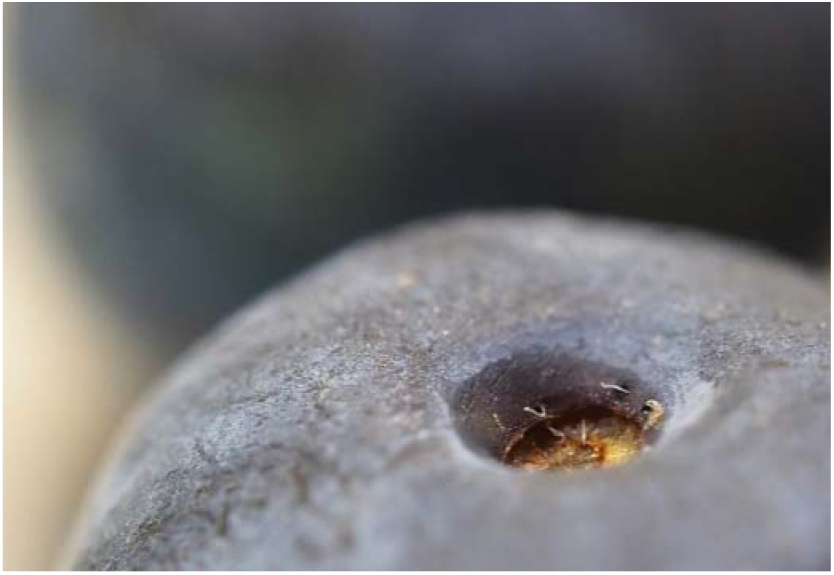
*D. suzukii* eggs are laid around the insertion of the fruit peduncle.

#### Maternal transmission

We used *D. melanogaster* population B, *D. suzukii* population B and the six microbial symbionts. Females were reared with males and associated with microbial symbionts five days before the experiment. Each experimental unit was constituted of one mature female and a wounded (*D. melanogaster*) or intact (*D. suzukii*) blueberry. Female and fruit were associated with a different microbial community (i.e. a different bacterium and a different yeast). For this experiment only, blueberries were inoculated with two different concentrations of microbial symbionts: a low concentration (5000 cells per microbial symbiont, decided in the light of previous estimates of cell numbers deposited by insects on fruit surfaces) and a high concentration (50 000 cells per microbial symbiont). Our initial goal was to test whether the concentration of fruit-associated microbial symbionts influences their transmission to the larvae, that was not the case (Table S1). A wet sterile cotton piece was added into each vial to ensure adult hydration. We created 36 female-fruit microbial combinations * two concentrations = 72 vials per *Drosophila* species. Females were disposed on fruits at 5 pm for 24 h. Controls without females were created to detect potential exogenous extracellular microorganisms. Adults were collected at the end of the day, crushed in PBS + 20% glycerol and stocked at −80°C. After five days, up to ten larvae were collected per fruit, pooled and crushed in PBS using a Tissue Lyser II. Right after crushing, larval samples were simultaneously plated on Galactose, Glucose, Mannitol (incubation at 24°C) and MRS plates (incubation at 30°C) to differentiate between and count microbial symbionts.

#### Male-mediated transmission

We used *D. melanogaster* population B, *D. suzukii* population B and the six microbial symbionts. The used adults were obtained from larvae associated with a combination of one yeast and one bacterium. Emerging male and female adults were kept five days in the same vials then separated per sex and re-associated with the original yeast + bacterium combination. Each experimental unit was constituted of one mature female, one mature male and a wounded (*D. melanogaster)* or intact (*D. suzukii)* blueberry. Prior to the experiment, each female, male and fruit were associated with a different microbial community (i.e. a different bacterium and a different yeast). For this experiment, blueberries were inoculated with 5000 microbial cells of each symbiont. A wet sterile cotton piece was added into each vial to ensure adult hydration. Per *Drosophila* species, we created 36 vials, one for each male-female-fruit microbial combination. In the early morning, the male was placed on the vial to enable the deposition of its microbial symbionts on the fruit surface. Note we verified the capability of males to deposit their symbionts on the fruit surface during a preliminary essay, this data will be presented in the next version of this work. To encourage the male to sit on the fruit, an axenic mature female kept in a small cage was added to the system (Figure 6). Male presence on the oviposition site was recorded eight times along the day. In the early morning of a second day, the captive axenic female was removed from the vial and the free-living gnotobiotic female was added. Male presence on the oviposition site was recorded eight times along this second day. Mating was also recorded every 30 min. Controls without females and males were created to detect potential exogenous extracellular microorganisms. Individuals were collected at the end of the second day, crushed in PBS + 20% glycerol and stocked at −80°C. After five days, up to ten larvae were collected per fruit, pooled and crushed in PBS using a Tissue Lyser II. Right after crushing, larval samples were simultaneously plated on Galactose, Glucose, Mannitol (incubation at 24°C) and MRS plates (incubation at 30°C) to differentiate between and count microbial symbionts.

**Figure 6.**
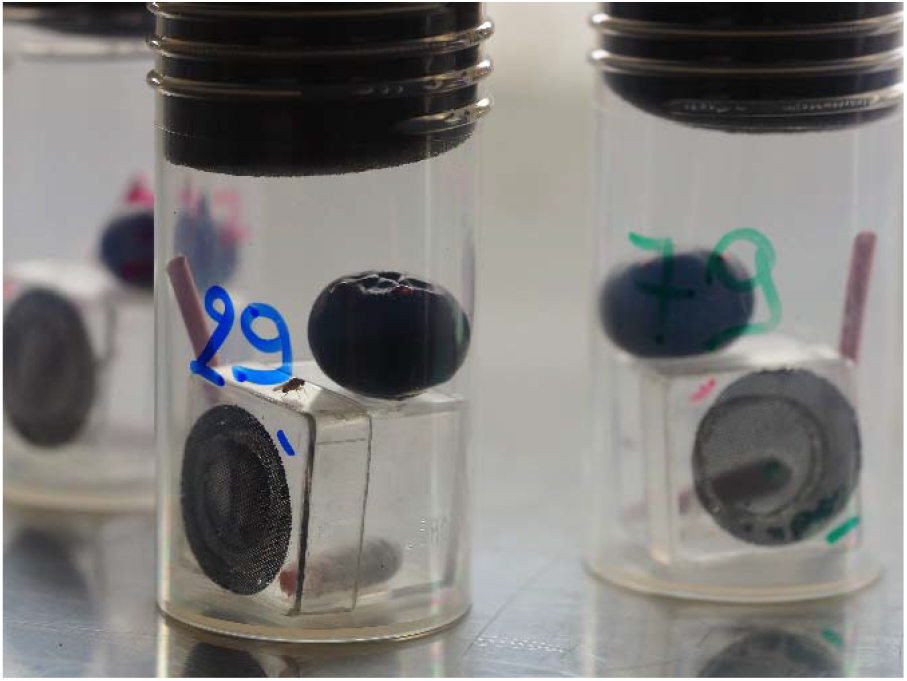
Example of experimental units used to test male transmission (on the first day, with a free-living gnotobiotic male and an axenic female in cage).

### Maintenance and transmission of microbial symbionts throughout the insect life cycle and between generations

We used *D. melanogaster* population B, *D. suzukii* population A and the three yeast isolates. Grape juice plates supplemented with the antifungal cycloheximide (1 μl/10 ml) were used to obtain yeast-free eggs from conventionally reared females. Eggs were deposited on surface-sterilized, incised grape berries (Behar et al., 2008) disposed on sterile vermiculite. After egg deposition, the wounds were inoculated with a single yeast strain (from overnight culture in YPD at 24°C). After the end of pupal formation, fruits were removed. Five freshly emerged adults of each sex were collected to evaluate yeast persistence through host metamorphosis. Other adults were placed in new experimental units. Each experimental unit was constituted of one male and one female that were reared with the same yeast strain. A petri dish with a wet cotton piece and sugar was disposed in the system to ensure fly survival. Right after setting of the system, a first grape berry inoculated with a second yeast strain was added. After five days, the fruit was removed and a second grape berry inoculated with a third yeast strain was added. Two days after, the fruit was removed and an incised, surface-sterilized grape berry was added to collect larvae. After one day, the adults were collected. Three days after, larvae were aseptically removed from fruit flesh. All adult and larval samples were crushed in PBS right after their collect using a Tissue Lyser II (Qiagen) and plated on Galactose and Glucose plates to differentiate and count yeast symbionts.

### Statistical analyses

GLM models with binomial distribution and logit function or poisson distribution and log function were used using JMP (SAS, 14.1). A backward stepwise model selection was used to eliminate non-significant terms from initial full models.

## Acknowledgements

We thank Laure Benoit, Marie-Pierre Chapuis and Romain Gallet for their help for the molecular identification of the microbial isolates used in this study and during the preliminary experiments.

