## Supplementary Materials for "The origin and maintenance of microbial symbionts in *Drosophila* larvae"

**Supplementary material 1.**

**Table S1.** Analysis of factors that may affected maternal transmission and environmental acquisition of microbial symbionts. Generalized linear models (GLM).

|  | **Maternal transmission**  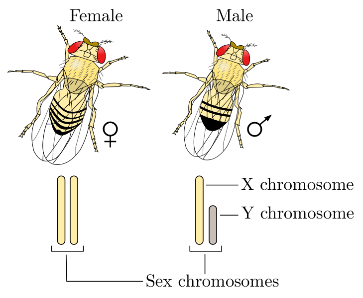 | **Environmental acquisition** |
| --- | --- | --- |
| Number of larvae collected | χ² = 4.85, df = 1, p = 0.0276 | χ² = 6.41, df = 1, p = 0.0113 |
| Fly species | p = 1, model could not compute | χ² = 1.98, df = 1, p = 0.1597 |
| Identity of the microbial symbiont of the female | χ² = 83.80, df = 4, p < 0.0001 | χ² = 5.00, df = 4, p = 0.2874 |
| Fly species * Identity of the microbial symbiont of the female | χ² = 11.76, df = 4, p = 0.0192 |  |
| Identity of the microbial symbiont of the fruit | χ² = 0.28, df = 4, p = 0.9910 | χ² = 44.38, df = 4, p < 0.0001 |
| Fly species * Identity of the microbial symbiont of the fruit |  | χ² = 4.26, df = 4, p = 0.3715 |
| Concentration of microbial symbiont of the fruit | χ² = 0.42, df = 1, p = 0.5149 | χ² = 0.25, df = 1, p = 0.6154 |

**Supplementary material 2. Presence of wild *D. melanogaster* and *suzukii* males and females in different fruits**

We evaluated *Drosophila* male and female presence rate on fruits. Wild *D. melanogaster* and *suzukii* mature adults were collected around Montpellier. For each species, six males and six females were placed on cages (n = 11) with a live strawberry plant and six strawberry fruits. During a day, we recorded number of individuals on fruits, plant and other parts of the cages every 30 min when flies are most active (Xuéreb A., Collet J. and Debelle A. pers. obs.), that is from 6 am to 10 am and from 6.30 pm to 9.30 pm (n = 16 observations). We tested whether the presence of individuals on fruits differed between the sexes and the fly species. We found that *Drosophila* males were more present on fruits compared to females for the two fly species (F_1,127_ = 10.56, p = 0.0012).

A	B

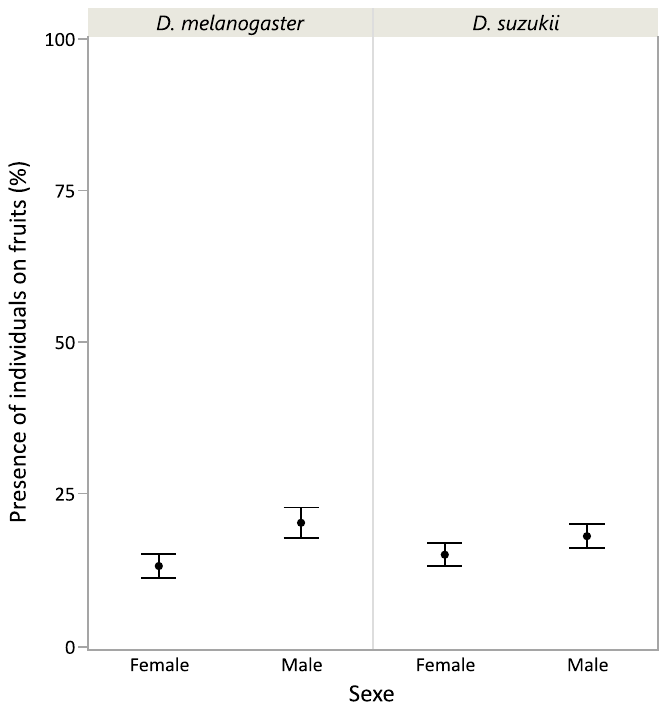

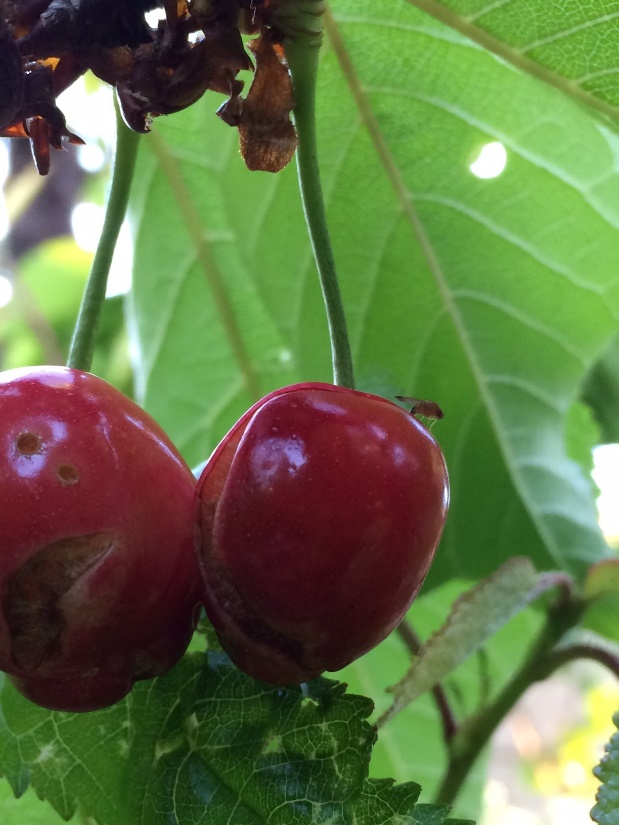

**Figure S2.** (A) *Drosophila* males were more present than females on fruits. Error bars show standard errors around the mean. (B) Observation of a *D. suzukii* male on a ripe cherry (photography: S. Fellous).

**Supplementary material 3.**

**Table S3.** Analysis of factors that may affected maternal transmission, male transmission and environmental acquisition of microbial symbionts. Generalized linear models (GLM).

|  | **Maternal transmission**  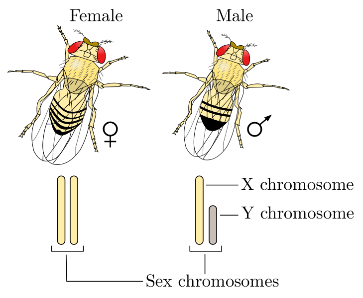 | **Male transmission** 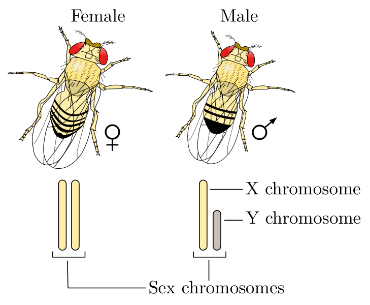 | **Environmental acquisition** |
| --- | --- | --- | --- |
| Number of larvae collected | χ² = 0.24, df = 1, p = 0.6253 | p = 1, model could not compute | χ² = 0.29, df = 1, p = 0.5893 |
| Fly species | χ² = 2.44, df = 1, p = 0.1186 | χ² = 12.10, df = 1, p = 0.0005 | χ² = 1.98, df = 1, p = 0.1591 |
| Identity of the microbial symbiont of the female | χ² = 48.57, df = 4, p < 0.0001 |  |  |
| Fly species * Identity of the microbial symbiont of the female | χ² = 4.46, df = 4, p = 0.3473 |  |  |
| Identity of the microbial symbiont of the male |  | χ² = 51.86, df = 4, p < 0.0001 |  |
| Fly species * Identity of the microbial symbiont of the male |  | χ² = 3.17, df = 4, p = 0.5301 |  |
| Male presence (%) |  | χ² = 0.35, df = 1, p = 0.5546 |  |
| Mating status |  | p = 1, model could not compute |  |
| Identity of the microbial symbiont of the fruit |  |  | χ² = 34.43, df = 4, p < 0.0001 |
| Fly species * Identity of the microbial symbiont of the fruit |  |  | χ² = 1.26, df = 4, p = 0.8678 |

**Supplementary material 4.**

**Table S4.** Analysis of factors that may influence presence of the different microbial symbionts in *Drosophila* adults. Generalized linear models (GLM).

|  | **Maintenance of larval symbionts** | **Acquisition of first environmental symbionts** | **Acquisition of second environmental symbionts** |
| --- | --- | --- | --- |
| Sex | χ² = 0.28, df = 1, p = 0.5976 | χ² = 2.50, df = 1, p = 0.1140 | χ² = 0.02, df = 1, p = 0.8902 |
| Fly species | p = 1, model could not compute | p = 1, model could not compute | χ² = 0.86, df = 1, p = 0.3527 |
| Identity of the larval symbiont | χ² = 6.53, df = 2, p = 0.0381 |  |  |
| Identity of the first environmental symbiont |  | χ² = 4.97, df = 2, p = 0.0834 |  |
| Identity of the second environmental symbiont |  |  | χ² = 23.54, df = 2, p < 0.0001 |
